## Supplementary material for "Linking mountaintop removal mining to water quality for imperiled species using satellite data": S1 Table

**S1 Table. 55 aquatic species that occur within Central Appalachia are listed on the Endangered Species List**.

| *Common name* | *Scientific name* | *ESA status (date listed)* | *Critical habitat* |
| --- | --- | --- | --- |
| Cumberland elktoe | *Alasmidonta atropurpurea* | Endangered (1/10/1997) | 8/31/2004 |
| Anthony's riversnail | *Athearnia anthonyi* | Endangered (4/15/1994) | - |
| Big Sandy crayfish | *Cambarus callainus* | Threatened (5/9/2016) | 1/28/2020^a^ |
| Guyandotte River crayfish | *Cambarus veteranus* | Endangered (5/9/2016) | 1/28/2020^a^ |
| Laurel dace | *Chrosomus saylori* | Endangered (9/8/2011) | 10/16/2012 |
| Diamond darter | *Crystallaria cincotta* | Endangered (8/26/2013) | 8/22/2013 |
| Spectaclecase | *Cumberlandia monodonta* | Endangered (4/12/2012) | - |
| Fanshell | *Cyprogenia stegaria* | Endangered (6/21/1990) | - |
| Dromedary pearlymussel | *Dromus dromas* | Endangered (6/14/1976) | - |
| Yellow lance | *Elliptio lanceolata* | Threatened (5/3/2018) | 2/6/2020^a^ |
| Cumberlandian combshell | *Epioblasma brevidens* | Endangered (1/10/1997) | 8/31/2004 |
| Oyster mussel | *Epioblasma capsaeformis* | Endangered (1/10/1997) | 8/31/2004 |
| Yellow blossom | *Epioblasma florentina florentina* | Endangered (6/14/1976) | - |
| Tan riffleshell | *Epioblasma florentina walkeri* | Endangered (9/26/1977) | - |
| Purple cat's paw | *Epioblasma obliquata obliquata* | Endangered (7/10/1990) | - |
| Green blossom | *Epioblasma torulosa gubernaculum* | Endangered (6/14/1976) | - |
| Northern riffleshell | *Epioblasma torulosa rangiana* | Endangered (1/22/1993) | - |
| Tubercled blossom | *Epioblasma torulosa torulosa* | Endangered (6/14/1976) | - |
| Snuffbox mussel | *Epioblasma triquetra* | Endangered (3/15/2012) | - |
| Turgid blossom | *Epioblasma turgidula* | Endangered (6/14/1976) | - |
| Spotfin chub | *Erimonax monachus* | Threatened (10/11/1977) | 9/22/1977 |
| Slender chub | *Erimystax cahni* | Threatened (10/11/1977) | 9/22/1977 |
| Bluemask darter | *Etheostoma akatulo* | Endangered (12/27/1993) | - |
| Duskytail darter | *Etheostoma percnurum* | Endangered (4/27/1993) | - |
| Kentucky arrow darter | *Etheostoma spilotum* | Threatened (11/4/2016) | 10/5/2016 |
| Cumberland darter | *Etheostoma susanae* | Endangered (9/8/2011) | 10/16/2012 |
| Shiny pigtoe | *Fusconaia cor* | Endangered (6/14/1976) | - |
| Finerayed pigtoe | *Fusconaia cuneolus* | Endangered (6/14/1976) | - |
| Cracking pearlymussel | *Hemistena lata* | Endangered (9/28/1989) | - |
| Pink mucket | *Lampsilis abrupta* | Endangered (6/14/1976) | - |
| Alabama lampmussel | *Lampsilis virescens* | Endangered (6/14/1976) | - |
| Birdwing pearlymussel | *Lemiox rimosus* | Endangered (6/14/1976) | - |
| Lee County cave isopod | *Lirceus usdagalun* | Endangered (11/20/1992) | - |
| Palezone shiner | *Notropis albizonatus* | Endangered (4/27/1993) | - |
| Yellowfin madtom | *Noturus flavipinnis* | Threatened (10/11/1977) | 9/22/1977 |
| Ring pink | *Obovaria retusa* | Endangered (9/29/1989) | - |
| Littlewing pearlymussel | *Pegias fabula* | Endangered (11/14/1988) | - |
| Snail darter | *Percina tanasi* | Threatened (11/10/1975) | 9/22/1977 |
| Blackside dace | *Phoxinus cumberlandensis* | Threatened (6/12/1987) | - |
| White wartyback | *Plethobasus cicatricosus* | Endangered (6/14/1976) | - |
| Orangefoot pimpleback | *Plethobasus cooperianus* | Endangered (6/14/1976) | - |
| Sheepnose mussel | *Plethobasus cyphyus* | Endangered (4/12/2012) | - |
| Clubshell | *Pleurobema clava* | Endangered (1/22/1993) | - |
| James spinymussel | *Pleurobema collina* | Endangered (7/22/1988) | - |
| Cumberland pigtoe | *Pleurobema gibberum* | Endangered (5/7/1991) | - |
| Rough pigtoe | *Pleurobema plenum* | Endangered (6/14/1976) | - |
| Slabside pearlymussel | *Pleuronaia dolabelloides* | Endangered (10/28/2013) | 9/26/2013 |
| Fluted kidneyshell | *Ptychobranchus subtentum* | Endangered (10/28/2013) | 9/26/2013 |
| Rabbitsfoot | *Quadrula cylindrica cylindrica* | Threatened (10/17/2013) | 4/30/2015 |
| Rough rabbitsfoot | *Quadrula cylindrica strigillata* | Endangered (1/10/1997) | 8/31/2004 |
| Cumberland monkeyface | *Quadrula intermedia* | Endangered (6/14/1976) | - |
| Appalachian monkeyface | *Quadrula sparsa* | Endangered (6/14/1976) | - |
| Rayed bean | *Villosa fabalis* | Endangered (3/15/2012) | - |
| Purple bean | *Villosa perpurpurea* | Endangered (1/10/1997) | 8/31/2004 |
| Cumberland bean | *Villosa trabalis* | Endangered (6/14/1976) | - |

Table shows the common and scientific names of listed species, dates on which species were added to the Endangered Species List and their current status (Threatened or Endangered), and the date at which any critical habitat was designated.

^a^Proposed critical habitat after completion of the study.
